# Developmental scarcity induces sex differences in social recognition and CA2 perineuronal nets

**DOI:** 10.64898/2026.01.09.698649

**Authors:** Renée C. Waters, Isha R. Gore, Casey J. Brown, Aimee L. Skweres-Gilmartin, Aarushi B. Rathaur, Elizabeth Gould

## Abstract

Early-life adversity has a lasting negative impact on social behavior in both humans and rodents. Sex differences exist in functional outcomes of postnatal stress, but underlying mechanisms remain incompletely explored. Using the limited bedding and nesting paradigm, a mouse model of developmental scarcity, we found sex differences in social behavior with adult males showing impaired social recognition and adult females showing intact social recognition with enhanced social novelty preference. Perineuronal nets, specialized extracellular matrix structures, are known to inhibit developmental plasticity and are concentrated in the CA2 region of the hippocampus in adulthood. We found higher intensity of CA2 perineuronal nets in adult males, but not adult females, after early life adversity. Focal degradation of the extracellular matrix in the CA2 region restored social recognition function in adult males previously subjected to early stress, suggesting that the effects of developmental adversity may be overcome by targeted intervention in adulthood.

## Introduction

In humans, childhood adversity increases susceptibility to several neuropsychiatric diseases in adulthood (Felitti et al., 1998). Among the neuropsychiatric diseases linked to early-life adversity (ELA), several present with deficits in social cognition, including posttraumatic stress disorder and major depressive disorder (Beck et al., 2009; Ladegaard et al., 2014; DiGangi et al., 2017). These neuropsychiatric diseases affect brain regions that are involved in sociocognitive processes, such as the hippocampus, amygdala, and prefrontal cortex (Tanimizu et al., 2017). Substantial evidence suggests that sex and type of ELA are important variables in determining neuropsychiatric outcomes in response to ELA. Emotional abuse is more commonly linked to alcohol use disorder in women while physical abuse and emotional neglect are more commonly linked to clinical depression and anxiety disorders in men (Prachason et al., 2024). Furthermore, several studies have raised the possibility that ELA can have differential neural consequences depending on sex. For example, emotional abuse is more closely tied to reduced hippocampal volume in males than in females, although the overall risk of psychopathology does not differ between sexes (Samplin et al., 2013).

Independent of neuropsychiatric diagnoses, ELA has been shown to affect social cognition. For example, children subjected to emotional abuse have a low threshold for assuming negative emotions on facial expressions in adulthood, while those subjected to emotional neglect have deficits in face recognition in adulthood (Iffland and Neuner, 2020). Given that social cognition impairments negatively affect quality of life (Schafer and Schiller, 2019), identifying neural mechanisms underlying ELA effects on social cognition is important for the goal of establishing points of intervention in adulthood. Toward this end, numerous studies have used animal models of ELA to explore neural and behavioral outcomes, with the majority focused on nonsocial behavioral outcomes, such as avoidance behavior, stress coping, and nonsocial cognition (Peña et al., 2017; Laham et al., 2022; Bolton et al., 2018; Yajima et al., 2018; Demaestri et al., 2024). One study showed that the ELA paradigm of limited bedding and nesting (LBN), a putative model of resource scarcity, reduced investigation time of novel versus familiar conspecifics (Kohl et al., 2015) although another study did not find any significant differences in this task (Shupe and Clinton, 2021). LBN has also been shown to reduce sociability (preference for social over nonsocial stimuli) in male, and not female, mice (Maulik et al., 2023).

The hippocampus, which plays an important role in social memory (Trinkler et al., 2009; Viskontas et al., 2009), undergoes considerable postnatal development, making it potentially vulnerable to the effects of early adverse events (Dick et al., 2022). Indeed, studies have shown that ELA reduces hippocampal volume in humans (Dannlowski et al., 2012; Paquola et al., 2016), and diminishes neurogenesis, dendritic arborization, and synaptogenesis in rodents (Mirescu et al., 2004; Leslie et al., 2011; Monroy et al., 2010). The CA2 region of the hippocampus has been referred to as a “social hub” because it integrates social information from multiple afferents to support social recognition (Hitti and Siegelbaum, 2014; Diethorn and Gould, 2023a). Afferents to this region, particularly from the dentate gyrus and supramammillary nucleus, undergo considerable development during the postnatal period (Diethorn and Gould, 2023b). In adulthood, the CA2 receives input from adult-born granule cells (abGCs) of the dentate gyrus, which have been linked to social recognition (Cope et al. 2020; Laham et al., 2024). The CA2 also contains a large concentration of perineuronal nets (PNNs), extracellular matrix structures implicated in plasticity inhibition, which have also been linked to social recognition in adulthood (Cope et al., 2022; Alexander et al., 2025). Furthermore, abGCs and PNNs are sensitive to early life stress (Mirescu et al., 2004; Catale et al., 2022; Laham and Gould, 2022), raising the possibility that ELA may influence social recognition by altering either or both plastic processes.

Here we show that exposure to the LBN paradigm impairs social recognition function in adult male, but not female, mice. We found that neither LBN males nor LBN females displayed differences in the number of abGCs in the dentate gyrus. However, LBN males, but not females, had increased PNN intensity, specifically of glycosaminoglycan (GAG) chains, in area CA2. Reduction of CA2 PNNs in adulthood restored social memory function in LBN males, showing that persistent dysfunction arising from early life stress can be corrected even after development has ended. These findings show that targeted intervention directed at ELA-specific mechanisms can restore social recognition function in adulthood.

## Methods

### Animals

All animal procedures were approved by the Princeton University Institutional Animal Care and Use Committee and were in accordance with the National Research Council Guide for the Care and Use of Laboratory Animals. Adult C57BL/6J mice were obtained from Jackson Laboratories and bred in the Princeton Neuroscience Institute. Initial studies used mixed-sex groups; however, follow-up investigations were male-only because LBN-induced behavioral impairments were not observed in females. All mouse pups were weaned from their dams on postnatal day (P) 21 and housed in same sex groups of 4-5 per cage. Mice were housed in Optimice cages on a reverse 12/12hr light/dark cycle.

### Limited Bedding and Nesting

Following cross-fostering on P2, pups were randomly assigned to the control-rearing or the LBN group. The LBN paradigm was adapted from previous studies (Gallo et al., 2019). Control litter cages containing standard bedding and a normal amount of nesting material were left undisturbed until weaning on P21. LBN pups and their dam were moved from their home cages and placed into a cage with a perforated metal floor, no bedding, and very little nesting material from P4-P11. Following P11, LBN mice were returned to cages with standard bedding and nesting until weaning on P21 (Figure 1A).

**Figure 1.**
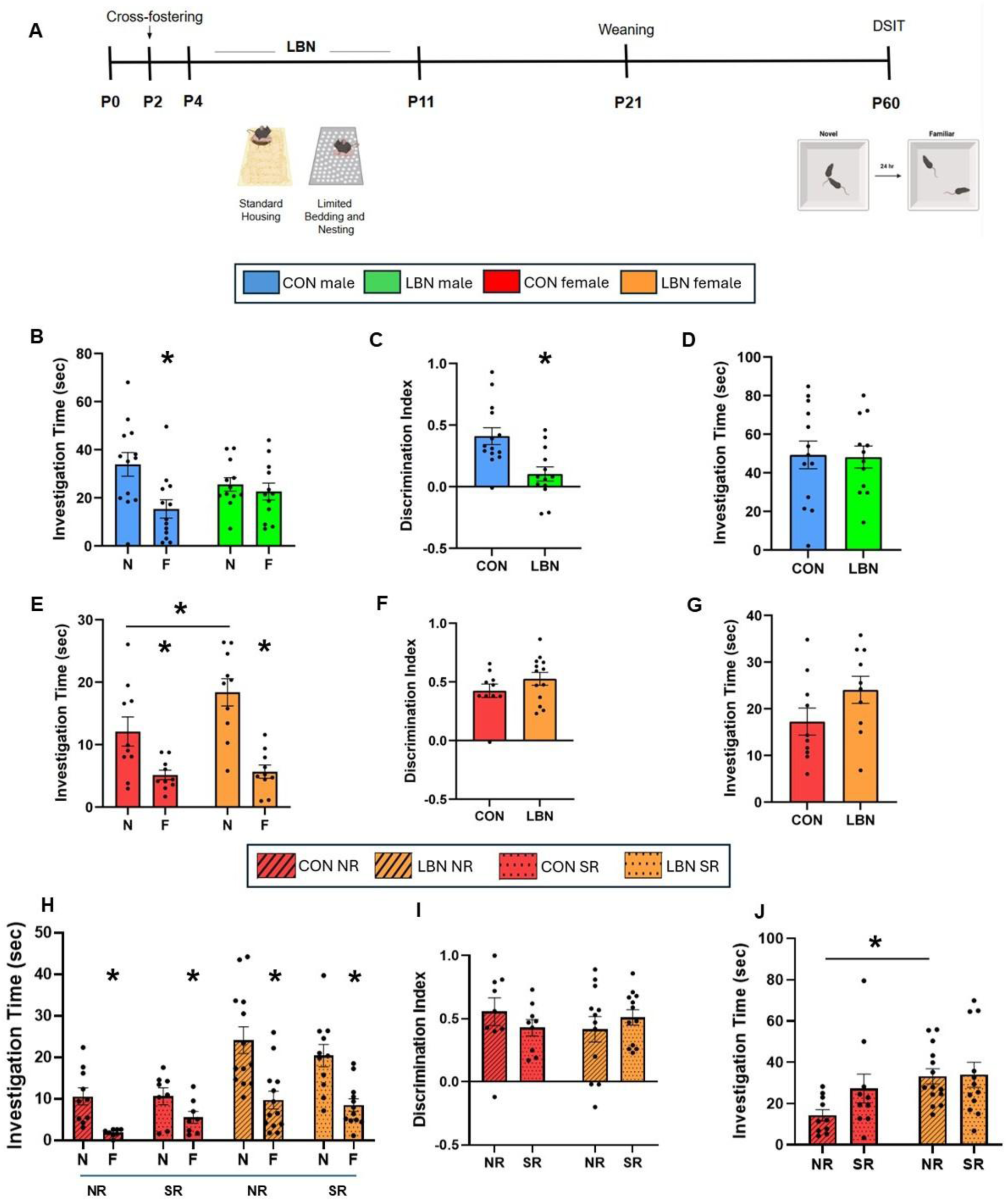
LBN impairs social discrimination in adult males, but not adult females. **A)** Experimental timeline and schematic of LBN paradigm and adult behavioral testing. **B)** Control-reared males displayed a significant reduction in investigation time between exposure to the novel mouse in trial 1 (N) and to the familiar mouse in trial 2 (F). LBN males showed similar investigation times in trial 1 and trial 2. **C)** LBN males had lower discrimination indices (N-F/N+F) than control-reared males. **D)** Overall investigation times (N+F) were not different between LBN and control-reared males. **E)** Both control-reared and LBN females display a significant reduction in time investigating the novel mouse in trial 1 to the familiar mouse in trial 2. LBN females show an increase in novel investigation compared to control-reared females. **F)** Control-reared and LBN females show similar discrimination indices. **G)** Overall investigation times (N+F) were not different between LBN and control-reared females. **H)** Control-reared and LBN females displayed lower investigation times with familiar than novel social stimuli regardless of whether the mice were tested when sexually receptive or nonreceptive. **I)** Control-reared and LBN females had similar discrimination indices regardless of whether they were tested when sexually receptive or nonreceptive. **J)** Control-reared females had lower overall investigation times than LBN females when they were nonreceptive, while sexually receptive control-reared and LBN females showed no statistical difference. *p < 0.05; N = novel social stimulus; F = familiar social stimulus; NR = nonreceptive; SR = sexually receptive

### Behavioral Analysis

On P60, the direct social interaction test (DSIT) was used to determine whether adult mice were able to discriminate between a novel and familiar mouse (Figure 1A) of the same sex and age as previously described (Cope et al., 2022; 2023). The DSIT was conducted in a 12-inch x 12-inch plexiglass arena with black walls and a clear base under low light (10-15 lux). Prior to the test beginning, mice were acclimated to the behavior testing room for at least 30 min and then habituated to the testing box for 5 min. Following habituations to the room and the testing arena, the mice were subjected to a 2-trial test. In trial 1 (novel), the test mouse was placed in the testing arena with a novel stimulus mouse. The mice were allowed to interact freely for 5 min. After a 24 hr delay in their respective home cages, trial 2 (familiar) began in which the test mouse was placed back in the testing arena with the same mouse for 5 min. The arena was thoroughly cleaned with 70% ethanol between each mouse and trial. Both trials were recorded using a single camera. A trained experimenter, blind to the treatment group, manually scored the time the test mouse spent investigating the stimulus mouse. Social investigation was defined as the test mouse directing its nose toward the stimulus mouse’s head, body, or anogenital region within 2 cm, following or initiating allogrooming. Because healthy adult mice prefer novelty, social recognition is inferred by a decrease in investigation time from the novel trial to the familiar trial (Moy et al., 2004). The discrimination index was calculated as follows: Discrimination Index = ([Novel-Familiar] / [Novel + Familiar]).

Because social behavior in females has been linked to sexual receptivity and estrous cycle stage (Chari et al., 2020; Sánchez-Andrade et al., 2011), female mice were subjected to vaginal smears for two weeks prior to behavioral testing, and their performance on direct social interaction was compared at stages of sexual receptivity (proestrus and estrus) with stages of sexual nonreceptivity (metestrus and diestrus).

### Surgical Manipulations

Adult male mice were deeply anesthetized with isoflurane (2-3%) and placed in a stereotaxic apparatus (Kopf) on a temperature-controlled thermal blanket for all surgeries. A microsyringe was used to deliver bilateral infusions of penicillinase (pnase Sigma Aldrich, Cat# P0389) or chondroitinase ABC (chABC Sigma Aldrich, Cat# C2905), an enzyme known to degrade GAG chains on PNNs (Bradbury et al., 2002; Pizzorusso et al., 2002) to the dorsal CA2 of control and LBN C57 adult males (15 nanoliter per injection at 3 nanoliters/min; AP: −1.82, ML: ±2.15, DV: −1.67). Mice were tested 10 days post infusion in the direct social interaction test. This post-surgical time point was selected because previous studies showed that at 5 days after CA2 chABC infusion, PNNs remain diminished and social recognition is impaired in control-reared mice whereas at 10 days after chABC infusion, CA2 PNNs are restored in control mice (Cope et al., 2022).

### Histology

Mice were anesthetized with Euthasol (Virbac) and transcardially perfused with cold 4% paraformaldehyde (PFA). Extracted brains were postfixed for 48 hr in 4% PFA at 4°C followed by an additional 48 hr in 30% sucrose at 4°C for cryoprotection before being frozen in tissue freezing medium at -80°C. Hippocampal coronal sections (40 μm) were collected using a cryostat (Leica). Sections were incubated for 1.5 hr at room temperature in a PBS solution that contained 0.3% Triton X-100 and 3% normal donkey serum. Sections were then incubated overnight while shaking at 4°C in the blocking solution that contained one or two of the following: mouse anti-3 microtubule-binding domain tau protein (3RTau, 1:500, Millipore, Cat# 05-803) to label immature neurons; rabbit anti-Purkinje cell protein 4 (PCP4, 1:500, Sigma-Aldrich, Cat# HPA005792) or mouse anti-regulator of G-protein signaling 14 (RGS14, 1:500, Antibodies Inc, Cat# 75-170) to label the CA2 region, rabbit anti-aggrecan (ACAN, 1:1000, Millipore, Cat# AB1031) to label the main neuronal chondroitin sulfate proteoglycan (CSPG) of PNNs or the plant lectin *Wisteria floribunda agglutinin* (WFA, 1:1000, Sigma Aldrich, Cat# L1516), to label GAG chains on PNNs.

For 3R-Tau immunohistochemistry, sections were subjected to an antigen retrieval protocol that involved incubation in sodium citrate and citric acid buffer for 30 min at 80°C before the blocking solution. All sections were then incubated for 1.5 hr at room temperature in secondary antibody solutions that contained combinations of the following secondaries: donkey anti-rabbit Alexa Fluor 568, donkey anti-mouse Alexa Fluor 488, donkey anti-rabbit Alexa Fluor 488, or Streptavidin 488 (1:500, Abcam). Sections were then counterstained with Hoechst 33342 for 10 min (1:5,000 in PBS, Molecular Probes), mounted onto slides, and cover slipped with Vectashield (Vector labs). Slides were coded until completion of the data analysis.

### Optical Intensity Measurements

Z-stack images of the dentate gyrus and CA2 were taken using a 40x objective with a 0.5 μm step size through the entire 40 μm tissue section on a Leica SP8 confocal with LAS X software (version 35.6). The CA2 region was defined by PCP4 or RGS14 labeling, which is specific to the CA2 in the hippocampal CA fields (San Antonio et al., 2014). Collected z-stack images of WFA and ACAN were analyzed for optical intensity in Fiji (NIH) as previously described (Cope et al., 2022). For overall intensity values in the CA2 region, the area, mean gray value, and integrated density of the pyramidal cell layer was calculated for each z-slice and the maximum integrated density value for each z-stack was taken from 3 sections per brain. Averages were obtained for each brain, and statistical comparisons were made between groups.

### Cell Density Measurements

The number of 3R-Tau+ cells was manually counted in the dorsal dentate gyrus of the hippocampus on 3 neuroanatomically matched sections using Fiji (ImageJ). The area measurements were collected using Fiji. The density of 3R-Tau was determined by dividing the total number of positively labeled cells by the volume of the subregion (ROI cross-sectional area multiplied by 40 μm section thickness).

### Statistical Analyses

Due to well-established sex differences in social investigation times (Karlsson et al., 2015; Granza et al., 2023), males and females were analyzed separately. For behavioral analyses involving two group comparisons, data sets were analyzed using either unpaired two-tailed Student’s t-tests or repeated measures two-way ANOVAs. For behavioral analyses involving chABC manipulations in adulthood, data sets were analyzed using a two-way ANOVA or a repeated measures three-way ANOVA. Tukey or Holm-Sidak post hoc comparisons were used to follow up any significant interactions or significant main effects that emerged from the ANOVAs. For histological analyses, LBN data were compared to controls for each sex with unpaired two-tailed Student’s t-tests. GraphPad Prism 9.2.0 (GraphPad Software) was used for statistical analyses and graph preparation.

## Results

### LBN impairs social recognition in adult male mice

Control-reared male mice had significantly lower investigation times for the familiar mouse than the novel mouse, while LBN male mice showed no significant differences in interaction time across trials (CON/LBN x novel/familiar interaction: F (1, 23) = 6.881, p = 0.0152; novel/familiar main effect: F (1, 23) = 13.09; p = 0.0014; novel/familiar post hoc comparisons: CON p = 0.0003; LBN p = 0.7472; Figure 1B). Social discrimination indices for LBN males were significantly lower than control-reared males (t_25_ = 3.418, p = 0.002; Figure 1C). LBN and control-reared male mice were similar in total social investigation times (novel+familiar) (t_23_ = 0.1205; p = 0.9051; Figure 1D), suggesting that the LBN-induced deficit in social recognition was not due to an overall change in social motivation but rather an inability to discriminate between novel and familiar mice.

### LBN enhances social novelty preference in female mice independent of sexual receptivity

Next, control-reared and LBN female mice were evaluated on the direct social interaction test using the same parameters. Like control-reared females, LBN females showed a significant decline in investigation time between novel and familiar trials (CON/LBN x novel/familiar interaction: F (1, 18) = 4.643; p = 0.0450; novel/familiar main effect: F (1, 18) = 54.20; p < 0.0001; novel/familiar post hoc comparisons: CON p = 0.0017; LBN p = 0.0001; Figure 1E). LBN females had comparable discrimination indices to control-reared females (t_21_=1.269; p=0.2184; Figure 1F). There was no statistical difference in overall investigation time (novel+familiar) between LBN and control-reared females (t_18_ = 1.660; p = 0.1143; Figure 1G), but LBN females had statistically higher investigation time of novel social stimuli (CON vs LBN novel: p = 0.0146; Figure 1E).

To determine whether stages of estrous/sexual receptivity interact with rearing condition, potentially obscuring negative effects of LBN on social recognition in females, we analyzed the data according to whether mice were in estrous stages associated with sexual receptivity (SR: proestrus and estrus) versus sexual nonreceptivity (NR: metestrus and diestrus). Although main effects were observed with rearing conditions (CON/LBN: F (1,12) = 22.09; p = 0.0005) and trial (novel/familiar: F (1,21) = 38.38; p = 0.0001), no effect of sexual receptivity stage was observed (F (1, 20) = 2.111; p = 0.1618; Figure 1H). Discrimination index data also showed no significant main effect of stage (NR/SR: F (1, 19) = 0.04185; p = 0.8401; Figure 1I). No overall differences in social investigation time (novel+ familiar) were detected across stages of sexual receptivity (F (1, 20) = 2.111; p = 0.1618) and LBN females only showed significantly higher investigation times during stages of nonreceptivity (CON vs LBN post hoc comparisons: NR p = 0.0233; SR p = 0.6245; Figure 1J).

### LBN does not affect the number of immature neurons in the dorsal dentate gyrus of males or females

abGCs are critical for social recognition in healthy mice (Cope et al., 2020), and through their afferents to area CA2, facilitate social memory retrieval and play an important role in overall CA2 network activity (Laham et al., 2024). Since LBN impaired social recognition in males, we hypothesized that a reduction in adult neurogenesis might contribute to this effect, so we investigated whether LBN affects the number of abGCs in males and females. Using 3R-Tau to label immature granule cells in dorsal dentate gyrus (Llorens-Martin et al., 2015), we found that 3R-Tau cell density was similar between LBN and control-reared mice with no statistical difference observed between sexes (t_22_ = 1.288, p = 0.2110; Figure 2A-D). This suggests that if adult neurogenesis plays a role in LBN-induced impairments in social recognition, it does not do so through detectable changes in the number of immature neurons.

**Figure 2.**
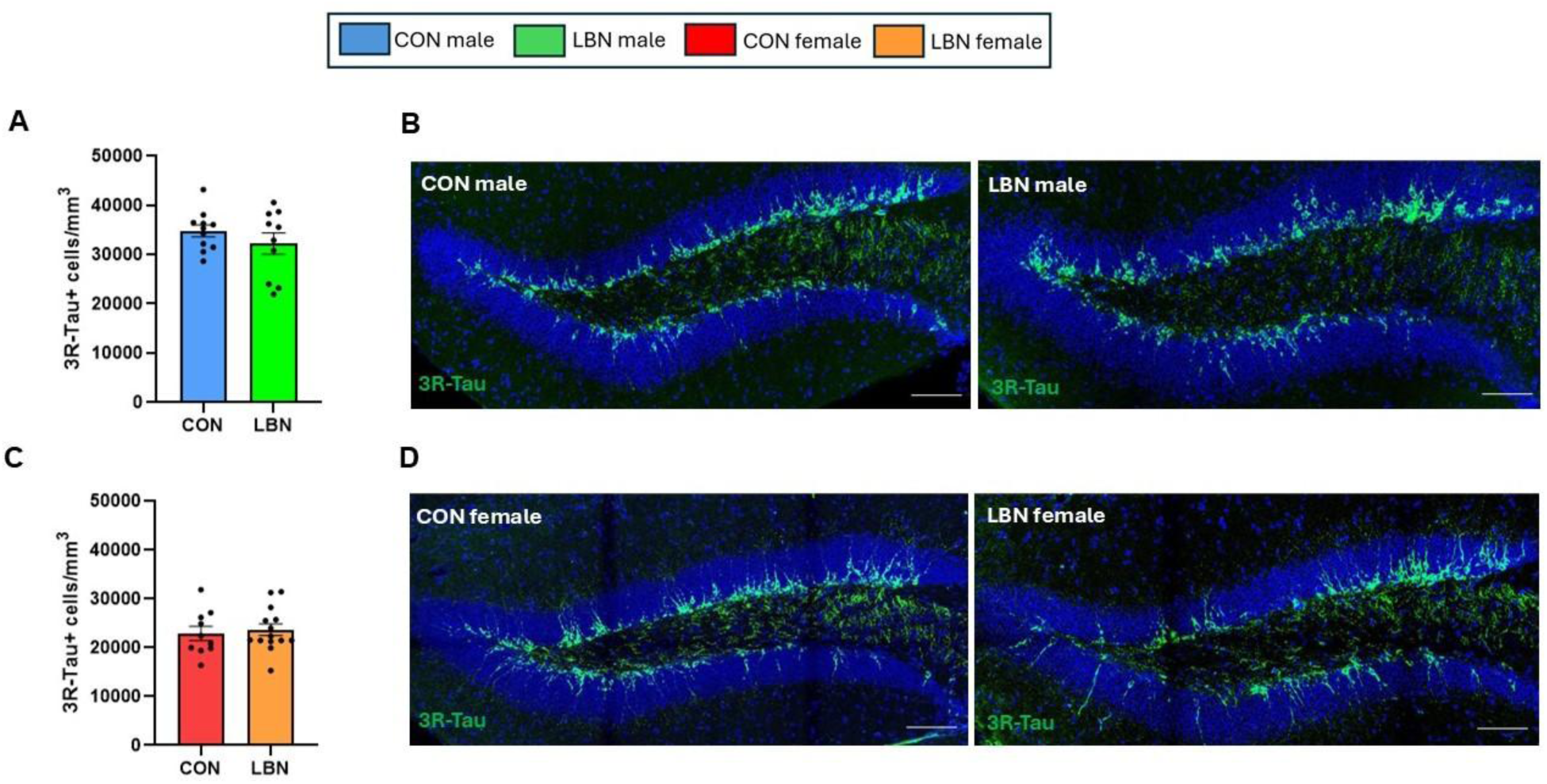
The number of immature neurons in the dorsal dentate gyrus is unaffected in adult male or female LBN mice. A, **C)** The number of neurons labeled with 3R-Tau in the dentate gyrus is similar between control-reared and LBN males and females. **B, D)** Representative confocal images of 3R-Tau+ cells (green) in dorsal dentate gyrus of control-reared and LBN adult males and females. Sections were counterstained with the DNA dye Hoechst 33342 (blue). Scale bars = 200 μm. 3R-Tau = 3 microtubule-binding domain tau protein.

### LBN male, but not female, mice exhibit increased CA2 PNN intensity compared to control-reared mice

Studies have shown that CA2 PNNs are important for the development and maintenance of social recognition abilities in mice (Diethorn et al., 2023b, 2025; Cope et al., 2022), so we investigated the possibility that CA2 PNNs differ between control-reared and LBN male mice. Labeling with the plant lectin *Wisteria floribunda agglutinin* (WFA), which binds to GAG chains on PNNs, showed a significant difference between groups. Confocal intensity analyses revealed that LBN male mice had >30% higher CA2 WFA intensity than control-reared male mice, an effect that was not observed in LBN female mice (t_18_ = 3.828, p=0.0012, Figure 3A-D). Despite finding increased WFA intensity in area CA2 of LBN male mice, we found no significant difference in intensity of the main neuronal chondroitin sulfate proteoglycan aggrecan (ACAN) between rearing conditions (t_19_ = 1.553, p = 0.1368; Figure 3E, F), suggesting that LBN males have altered PNN composition (more GAG chains) as opposed to an overall increase in all PNN components. No differences were observed in ACAN intensity in area CA2 between control-reared and LBN females (t_17_ = 0.2558; p = 0.8012; Figure 3G,H). Furthermore, no LBN-induced differences were observed in the cross-sectional area of the CA2 region in males (Control: 35625 ± 2430, LBN: 36059 ± 2614; t_20_ = 0.1197; p = 0.9059) or females (Control: 32930 ± 2665, LBN: 32772 ± 2406; t_18_ = 0.04387; p = 0.9655).

**Figure 3.**
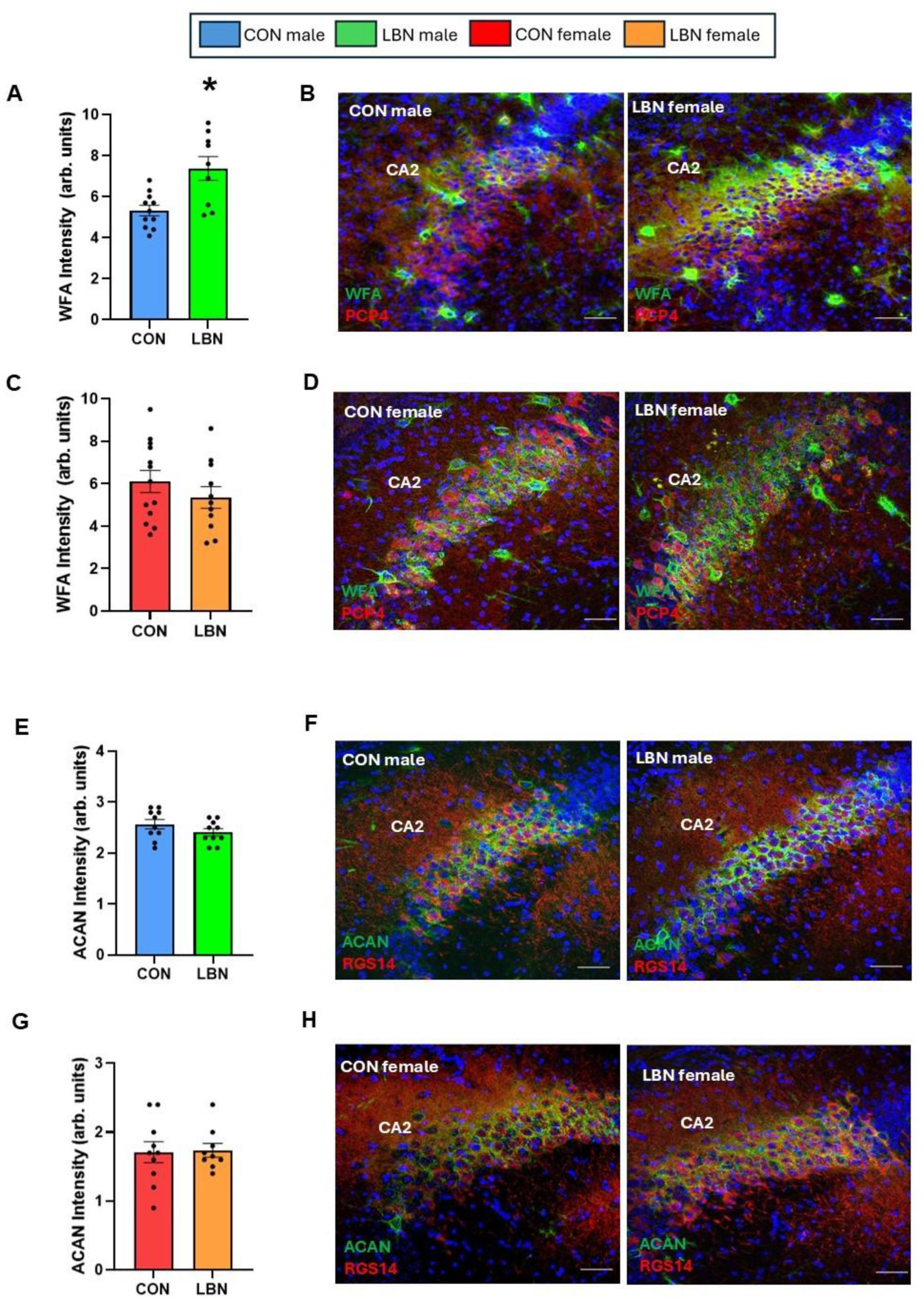
LBN males have increased WFA+ but not ACAN+ intensity in the CA2 region. **A)** LBN males had significantly higher intensity of WFA+ staining in the pyramidal layer of the CA2 compared to control-reared males. **C)** This effect was not observed in females, which showed no difference between rearing groups. **B, D)** Confocal images of WFA+ staining (green) in dorsal CA2 labeled with PCP4 (red) of control-reared and LBN males and females. **E, G)** LBN males and females showed no difference in ACAN+ intensity in the pyramidal cell layer of the CA2 compared to control-reared males and females respectively. **F, H)** Confocal images of ACAN labeling (green) in dorsal CA2 labeled by RGS14 (red) of control-reared and LBN males and females. Sections were counterstained with the DNA dye Hoechst 33342 (blue). Scale bars = 50 μm; *p < 0.05. WFA = *Wisteria floribunda agglutinin*; ACAN = aggrecan; PCP4 = Purkinje cell protein-4; RGS14 = regulator of G-protein signaling 14.

### Reducing CA2 PNNs in LBN males restores social discrimination

Because higher CA2 WFA intensity paralleled social recognition impairment in LBN males, we next investigated whether reducing GAG chains on PNNs with the enzyme chondroitinase ABC (chABC) (Bradbury et al., 2002; Pizzorusso et al., 2002) in the CA2 would improve social memory. Control-reared and LBN adult males were infused with chABC or the control enzyme penicillinase (pnase) in the bilateral CA2 (Figure 4A). Mice were tested on the direct social interaction task 10 days post-injection, a timepoint when WFA-labeled PNNs in the CA2 of control mice have regrown after chABC infusion to control-reared baseline levels and there is no longer a social recognition deficit (Cope et al., 2022). There was a significant interaction between rearing condition and CA2 infusion treatment in both trial-by-trial (novel, familiar) analysis (CON/LBN x chABC/pnase interaction: F (1, 39) = 4.529, p = 0.0397; Figure 4B) and the discrimination index (CON/LBN x chABC/pnase interaction F (1,43) =38.85, p < 0.0001; Figure 4C) As expected based on previous findings (Cope et al., 2022), we found that 10 days after surgery, control-reared males with pnase and chABC CA2 infusions had typical social recognition abilities, investigating novel mice significantly more than familiar mice (Figure 4B, C). By contrast, LBN males with pnase infusions showed impaired social recognition, investigating novel and familiar mice at similar levels (Figure 4B, C). However, 10 days after chABC LBN males had restored social discrimination abilities comparable to pnase control-reared males (Figure 4B, C). LBN mice given pnase had significantly lower discrimination indices compared to all other groups (Figure 4C). After perfusion, the efficacy of chABC to reduce PNNs in LBN males was assessed with WFA labeling. LBN males treated with chABC had significantly lower WFA intensity than LBN males treated with pnase (t_14_ = 2.171, p = 0.0477; Figure 4D, E).

**Figure 4.**
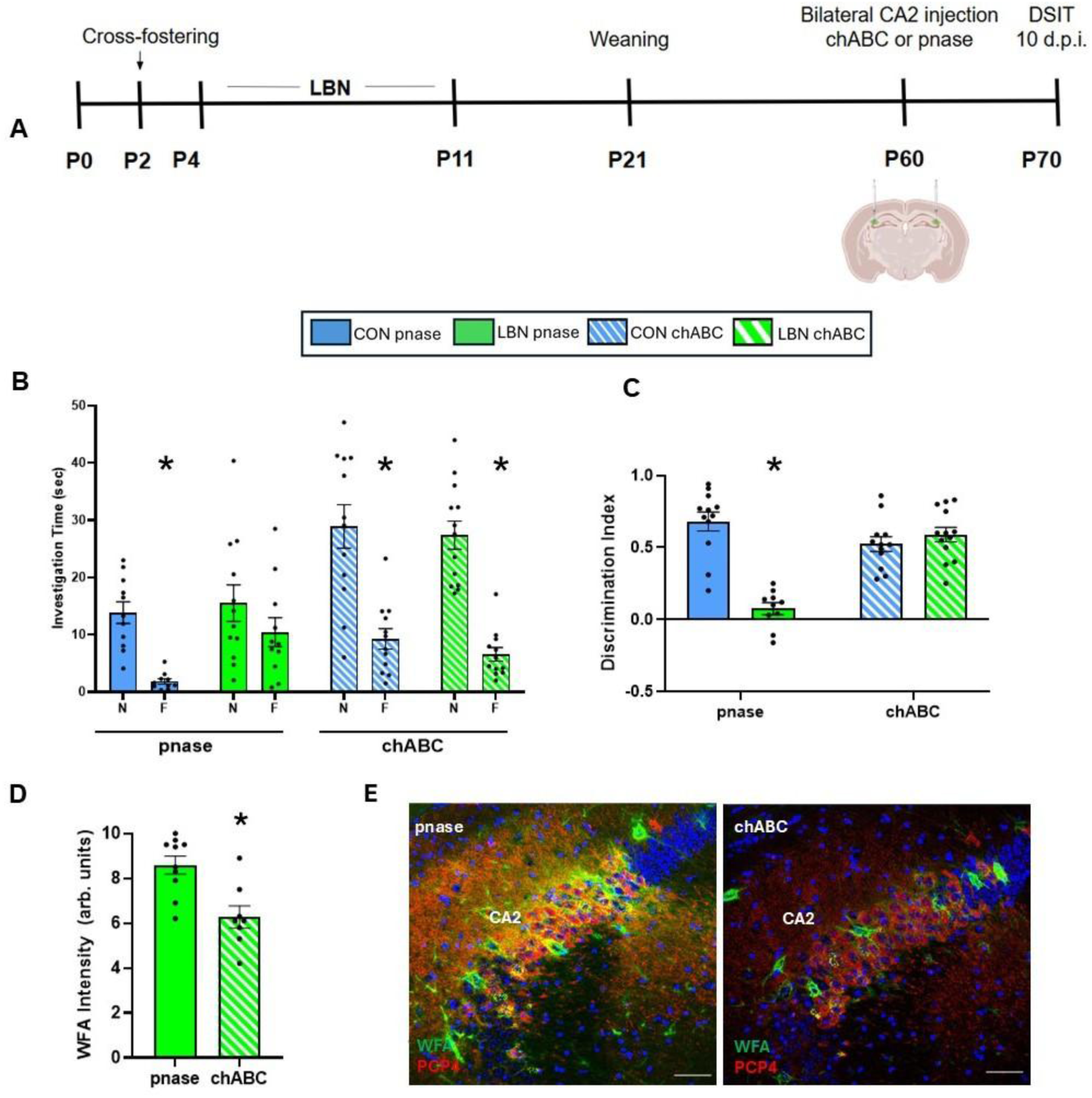
Reducing CA2 PNNs restores social recognition function in LBN males. **A)** Timeline of LBN chABC behavioral experiment. **B)** Reducing CA2 PNNs levels restored social discrimination function in LBN males. Control pnase, control chABC, and LBN chABC males display a significant reduction in time investigating the novel mouse (N) in trial 1 to the familiar mouse (F) in trial 2. LBN pnase males showed similar investigation times in trial 1 and trial 2. **C)** LBN pnase males had significantly lower discrimination indices compared to control pnase, control chABC, and LBN chABC males. **D)** LBN chABC males had lower WFA intensity in the CA2 than LBN pnase males. **E)** Confocal images of WFA labeling (green) in dorsal CA2 labeled with PCP4 (red) of LBN pnase and LBN chABC males. Sections were counterstained with the DNA dye Hoechst 33342 (blue). Scale bars = 50 μm; *p < 0.05. pnase = penicillinase; chABC = chondroitinase ABC; WFA = *Wisteria floribunda agglutinin*; PCP4 = Purkinje cell protein 4; DSIT = direct social interaction test; dpi = days post-injection

## Discussion

We found that LBN impaired social recognition function in adult males. In contrast, we found that LBN did not impair social recognition function in females but increased novel social investigation. In females, protection from LBN-induced social impairment occurred regardless of sexual receptivity/stage of estrous. We also found sex differences in measures of hippocampal plasticity. We found no differences in number of immature neurons in the DG of males or females after exposure to LBN. However, LBN-reared males had significantly higher PNN intensity in the CA2 compared to controls, while females exposed to LBN showed no differences, Because LBN-induced social memory impairment was associated with atypically high PNNs in adult males, we tested the effects of lowering PNNs in the CA2 region in adulthood and found improved social recognition. These findings suggest that adult intervention may be sufficient to temporarily restore some negative effects of developmental adversity.

### Excessive CA2 PNNs inhibit social recognition in LBN males

Previous studies have shown that the CA2 region plays an important role in social recognition both during development (Laham et al., 2022; Diethorn and Gould, 2023a) and in adulthood (Hitti and Siegelbaum, 2014; Laham et al., 2022; 2024). There is an unusually high concentration of PNNs in the CA2, which surround both inhibitory interneurons, as is typical for other hippocampal subregions, and pyramidal cells (Carstens et al., 2016; Cope et al., 2022). The emergence of PNNs in the CA2 appears to coincide with social recognition function during development (Diethorn and Gould, 2023a). In adulthood, CA2 PNNs are important for the ability to distinguish between novel and familiar mice; degradation of PNNs impairs this function in otherwise healthy mice (Cope et al., 2022). Conversely, inbred and genetic mouse models of social dysfunction have been associated with excessive PNNs in the CA2; some functional deficits can be restored by lowering levels of PNNs to control values (Carstens et al., 2021; Cope et al., 2022; Diethorn et al., 2025). These findings are consistent with our data showing a restoration of social recognition function in LBN males after reduction of PNNs by local infusion of the degradative enzyme chABC into the CA2 region. Furthermore, these findings suggest that the influence of LBN on circuitry underlying social recognition is not permanent and instead, can be repaired (at least temporarily) by adult intervention. The duration of these improvements, as well as the possibility of permanently restoring function, remains unknown.

Our results suggest that the influence of LBN on CA2 PNNs in adult males may be specific to GAG chains, as WFA intensity was increased but no effects were observed with the main neuronal CSPG ACAN. WFA labeling and ACAN immunostaining often co-label PNNs in the hippocampus, but there are reports of PNNs labeled by one marker and not the other (Yamada and Jinno, 2016; Laham et al., 2022). Furthermore, other mouse models of pathological states, such as neuropathic pain and Alzheimer’s Disease, have been associated with changes in one of these markers and not the other (Scarlett et al., 2022; Tansley et al.2022; de Vries et al., 2025), suggesting that maintaining an optimal balance of PNN constituents may be important for healthy function. Future studies should investigate whether enzymes associated with increased glycosylation of CSPGs are altered in adult LBN males and how these changes may affect social recognition.

PNNs appear to restrict plasticity by limiting postsynaptic sites available for synaptogenesis and/or by concentrating inhibitory neurotransmitter receptors so that inhibition is more robust (Wingert and Sorg, 2021). Previous work in other systems has shown that PNNs are sensitive to the environment (Laham and Gould, 2022) and neuronal activity, the latter of which appears to be negatively correlated with CA2 PNN intensity (Carstens et al., 2016; 2021). Thus, it is possible that the male LBN CA2 has higher PNNs because neuronal activity is diminished in the region. There are many ways this could occur, including by LBN affecting intrinsic properties of CA2 excitatory or inhibitory neurons, which have been shown to be sensitive to PNN degradation (Carstens et al., 2021). Another possibility is that LBN might alter developmental innervation from afferents outside of the hippocampus, the majority of which are excitatory (Diethorn and Gould, 2023a). If LBN diminishes excitatory inputs to the CA2, the corresponding decrease in activity in the postsynaptic region might contribute to excessive PNN formation. While the mechanism by which increased PNN intensity in the CA2 of LBN males impairs social recognition remains unknown, our data suggest that the effect may be reparable with interventions designed to correct PNN levels in adulthood.

### Sex differences in LBN effects on social recognition

Our results demonstrate divergent behavioral and cellular effects of LBN in males and females. In adulthood, LBN males have impaired social recognition ability. In striking contrast, adult female LBN mice exhibit robust social recognition abilities, with higher social novelty responses compared to control-reared females. These findings raise questions about whether the sex differences arise from differential developmental experience or whether they emerge over time. Regarding the former possibility, a few studies have shown that rodent mothers display differential care toward male and female pups, perhaps due to a greater need for licking and grooming to stimulate micturition in males (Sakamoto et al., 2021; Maggi et al., 1986). Thus, LBN, which is known to alter maternal behavior (Gallo et al., 2019; Shupe and Clinton, 2022; Baracz et al., 2020; Bailoo et al., 2015), may induce differential experiences in male and female pups, with greater long-term effects observed in males. However, it remains possible that the effects of LBN are largely similar on male and female pups, but that the females recover as time passes, possibly due to changes during puberty. Along these lines, we tested the LBN adult females during sexually receptive and sexually non-receptive stages of the estrous cycle and found no differences in social behaviors suggesting that cyclic changes in hormone levels are not sufficient to reveal dysfunction. Thus, future studies should focus on understanding how changes in signaling molecules during the peripubertal period, or differential expression of stress-related genes, in females interact with circuitry underlying social recognition.

## Acknowledgments

The authors thank Hunter Worth and Sharon Powley for technical assistance. This work was supported by NIMH R01MH117459 to EG and NSF GRFP 2021318039 to RCW.

